# Net-shaped DNA nanostructure designed for rapid/sensitive detection and potential inhibition of SARS-CoV-2 virus

**DOI:** 10.1101/2022.05.04.490692

**Authors:** Neha Chauhan, Yanyu Xiong, Shaokang Ren, Abhisek Dwivedy, Nicholas Magazine, Lifeng Zhou, Xiaohe Lin, Tianyi Zhang, Brian T. Cunningham, Sherwood Yao, Weishan Huang, Xing Wang

**Author notes:** A U.S. provisional patent has been filed in November 2020 based on part of the study reported in this manuscript.

## Abstract

We present a net-shaped DNA nanostructure (called “DNA Net” herein) design strategy for selective recognition and high-affinity capture of the intact SARS-CoV-2 virions through spatial pattern-matching and multivalent interactions between the aptamers (targeting wild type spike-RBD) positioned on the DNA Net and the trimeric spike glycoproteins displayed on the viral outer surface. Carrying a designer nanoswitch, the DNA Net-aptamers releases fluorescent signal upon virus binding that is easily read by a hand-held fluorimeter for a rapid (in 10 mins), simple (mix-and-read), sensitive (PCR equivalent), room temperature compatible, and inexpensive (∼ $1.26/test) COVID-19 test assay. The DNA Net-aptamers also impede authentic wild-type SARS-CoV-2 infection in cell culture with a near 1×10^3^-fold enhancement of the monomeric aptamer. Furthermore, our DNA Net design principle and strategy can be customized to tackle other life-threatening and economically influential viruses like influenza and HIV, whose surfaces carry class-I viral envelope glycoproteins like the SARS-CoV-2 spikes in trimeric forms.

## INTRODUCTION

COVID-19, caused by the SARS-CoV-2 virus, has resulted in a pandemic responsible for severe illness and ∼ 2.53 deaths by early March 2021^1^. A key failure observed during the early COVID-19 pandemic in the health systems across the world has been the inability to provide rapid and accurate diagnosis using the existing paradigms of nucleic acid and antigen testing. The limitations of the most widely used SARS-CoV-2 diagnostic technologies in early days of the pandemic can be attributed to the limited availability of valid test kits, the availability and testing bandwidth of certified testing facilities, and a lengthy and expensive laboratory procedure to obtain a result and provide diagnostic information to the patients^2,3^. The time and expense associated with the currently dominant tests for virus detection stem from the requirement for detecting nucleic acid sequences of one pathogen that generally necessitate the pre-processing of samples and amplification of the target sequences. In order to perform these genome-based tests, stringent and technically challenging laboratory protocols for virus particle lysis, RNA extraction from RNA viruses, RNA reverse transcription (RT), and enzymatic amplification of specific nucleic acid sequences by polymerase chain reaction (PCR) or alternatives such as loop-mediated isothermal amplification (LAMP) are a prerequisite^4^. Although such methods can be automated and performed with a high throughput using sophisticated equipment, all the nucleic acid test (NAT) methods require complex chemistries, accurate temperature control, enzymes, and its conditional buffer solutions, and many sample-handling steps.

Furthermore, even when PCR tests are performed correctly, the assays detect the presence of SARS-CoV-2 genetic materials rather than the presence of virions that are actively infective. A growing body of scientific and medical reports suggest that SARS-CoV-2 PCR positivity could be a result of residual viral RNA or dead/inactive viruses. Therefore, a positive PCR test cannot accurately determine the presence of active and infectious viruses in the tested individuals^5,6^. As an alternative to NAT, antigen-based tests can be rapid and portable for detecting the presence of infectious viral entities. However, rapid antigen tests suffer from low sensitivity, which renders these incapable of detecting low viral loads during early infection.

There is an urgent need to improve the performance of virus diagnostics both in point-of-care (POC) settings and with high sensitivities to help curb the spread of highly contagious diseases like COVID-19^7-9^. Nanomaterials and nanotechnology have provided promising solutions to tackling infectious diseases^10-19^. We have previously demonstrated a “DNA Star” strategy for creating a biosensor by targeting the immobile envelope protein clusters (called ED3) exposed on the dengue virus (DENV) outer surface^16^. DENV envelope proteins are rigid and arranged into a shell-like structure on the viral surface, making them suitable for a star-shaped rigid matching. In contrast, class I viral envelope glycoproteins on membraned viruses such as coronaviruses, influenza, and HIV, have flexible stems and mobile roots, presenting a huge challenge for pattern-matching, which cannot be tackled by the “DNA Star”, or similar strategy designed for targeting rigid surface antigens. Thus, a change in strategy that would allow for higher degrees of flexibility for virus-sensor interactions is vital for COVID-19 diagnosis using DNA nanotechnology.

**Scheme 1.**
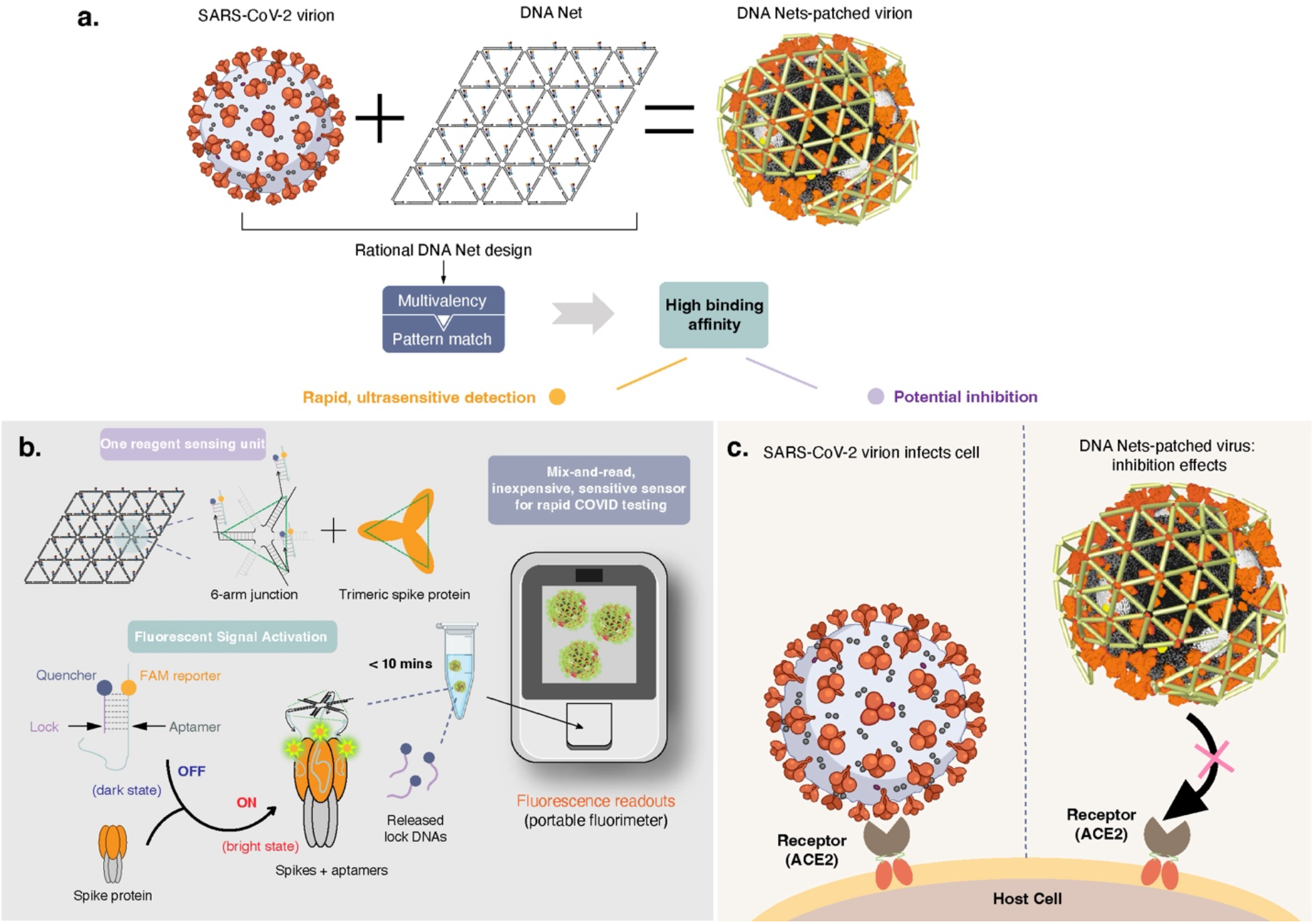
Schematic of Viral Capture and Reading/inhibition (VCRi). **a**, Rationally designed DNA Net vertices incorporate WT spike RBD-targeting aptamers to form an array of trimeric clusters that each has a mechanical match to the intra-spacing of the protomers within the trimeric spike protein on the SARS-CoV-2 viral surface. DNA Net-aptamers can enable dynamic spike clustering, resulting in multiple sites of high-affinity attachment. **b**, The sensing motif in the DNA Net sensor comprises of a FAM tagged aptamer quenched by a partially complementary ‘Lock’ DNA. The interaction of the aptamers on the DNA Net sensor with the virus triggers a rapid release of multiple Lock DNA to unquench the FAM reporters even at a low viral concentration (*high detection sensitivity*). The fluorescent output can be read by a portable fluorimeter. **c**, The DNA Net-aptamers complex patched on a virion blocks spike-ACE2 interactions on a host cell surface to inhibit virus infection (*high neutralization potency*).

In this work, we present a “three-layer” Designer DNA Nanostructure (DDN) design principle/strategy for selective recognition and high affinity capture of intact SARS-CoV-2 virions. The DDN interacts via pattern-matching and multivalent interactions among the aptamers on the DNA Net and the trimeric spikes displayed on the viral surface. The aptamer switches patterned on the DDN are engineered to spontaneously release fluorescent signal upon detecting SARS-CoV-2 virus. Specifically, in this approach, called Viral Capture and Reading/inhibition (VCRi, **Scheme 1**), a net-shaped DDN, also referred as DNA Net henceforth, is rationally designed and subsequently synthesized to organize multiple aptamers that were selected to target the wild type (WT) SARS-CoV-2 spike protein’s receptor binding domain (RBD) via SELEX (DNA Net-aptamers, henceforth)^20^. Aptamers are patterned on the DNA Net into an array of trimeric clusters (or called “tri-aptamers” henceforth), each of which has a precise match to the intra-spacing of the trimeric spike proteins. Since the spikes are mobile on the SARS-CoV-2 viral outer surface, the DNA Net-aptamers can potentially enable a dynamic clustering of spikes to provide a maximal spike packing density. Similar to DNA scaffolds guided clustering of cell surface mobile proteins^21-24^, clustering of spikes using DNA Net can achieve a global tri-aptamer-spikes pattern matching by correcting deviations of inter-spike spacing through the multiple sites of attachment, resulting in an enhanced virus binding affinity (**Scheme 1a**). To transform the DNA Net-aptamers into a virus sensor, an aptamer nanoswitch was designed and introduced for robust fluorescent signal release, in which the SARS-CoV-2 spike-specific binding aptamer was tagged with a fluorescent reporter, along with a quencher-labeled “lock” DNA that forms a partial duplex with the aptamer. These aptamer-quenched fluorophore units were then organized into a trimeric cluster on each knot/vertex of the DNA Net (**Scheme 1b**). Like a fishing net in water, the DNA Net has mechanical bendability to form a concave shape in solution^16,25,26^, also promoted by the unpaired thymine bases (Ts) present on each knot of the DNA Net in our design^27^. When exposed to SARS-CoV-2 virions, the bendable DNA Net can curve itself to match the radius of curvature of viral particle surface, promoting multivalent and pattern-matching interactions between the patterned aptamer-lock pairs and the clustered spike proteins. This strong, rapid, and selective interaction offered by DNA Net-aptamers then greatly promotes the departure of the lock DNA from the aptamer, in turn separating the fluorophore from the quencher. A fluorescent signal is released within 10 minutes of mixing the DNA Net sensor reagent with the test sample, through a single-step, single reagent, room temperature workflow. The resulting fluorescence is easily detected by a portable fluorimeter^28^, achieving a limit of detection (LoD) of 1,000 viral genome copies/mL in the artificial saliva-containing solution. Such fluorescent signals can also be read by a high throughput platform such as a qPCR system in laboratory settings^16^. Additionally, the DNA Net-aptamers exhibited inhibition of SARS-CoV-2 infection in cell culture with a nearly 1×10^3^-fold enhancement over the monomeric aptamer, suggestive of a significant potential of our DNA Net as a candidate therapeutic agent for viral inhibition by blocking virion-host cell interactions (**Scheme 1c**).

## RESULTS

### DNA Net-aptamers design strategy

For SARS-CoV-2, the structure of the trimeric spike cluster has been elucidated using cryogenic electron microscopy (cryo-EM)^29,30^. The diameter of the SARS-CoV-2 virion is ∼ 100 nm (ref.^31^). The intra-molecular spacing between the aptamer binding sites on RBDs on the spike protomers of a given trimer cluster in the closed, RBD “down” prefusion conformation is ∼ 6 nm (**Fig. 1a, top view**). Previous studies show that approximately 60% of the spike trimers on the surface of the SARS-CoV-2 virion are in the closed prefusion conformation, and the remaining trimers each have one spike RBD in an open, “up” prefusion conformation with a slightly smaller spacing to the adjacent RBDs in a trimer cluster^32^. The spike trimers are in a constant state of motion owing to the fluid nature of the viral membrane. The virion size, surface curvature, surface spike structure, and membrane fluidity of the SARS-CoV-2 virus are largely consistent with the SARS-CoV-1 virus that contains 50-100 spike trimers/virion with an inter-spike trimer spacing of 15 nm (Ref.^33,34^). However, the experimentally determined density and distribution of spike proteins on SARS-CoV-2 surface varies across different studies^31,32,35,36^. This is likely due to the damage and/or loss of spike proteins from the viral surface during sample preparation steps involved in these studies (ultracentrifugation, sample resuspension, flash freeze, etc.)^32,35,36^. Based on the structure of SARS-CoV-2 trimeric spike protein^29,30^, the minimum possible distance between two neighboring spike trimers without any steric hindrances is estimated to be ∼14-15 nm. Thus, the DNA Nets were designed and synthesized from macromolecular net-shaped DNA nano-scaffolds with triangular units to pattern spike-RBD-targeting aptamers into an array of tri-aptamer clusters with a ∼ 6 nm intra- and ∼ 15 nm inter-trimer spacing to detect intact virions for rapid, sensitive COVID testing, and potent viral inhibition. To capture multiple spike trimers that move into the DNA Net vicinity, the triangular tile-based DNA Net architecture provides ample flexibility and maximum coverage to cluster spike proteins on spherical viral surface^37^. While the 6 nm intra-trimer spacing provides a first layer pattern-matching based aptamer-spike interactions, we speculated and experimentally validated herein that the 15 nm inter-tri-aptamer spacing on the DNA Net allows for high spike clustering efficiency that increases the number of spike trimers that can simultaneously interact with the tri-aptamers on the DNA Net. The resultant pattern-matching, multivalent based interactions yield a rapid virus binding with high avidity.

**Fig. 1.**
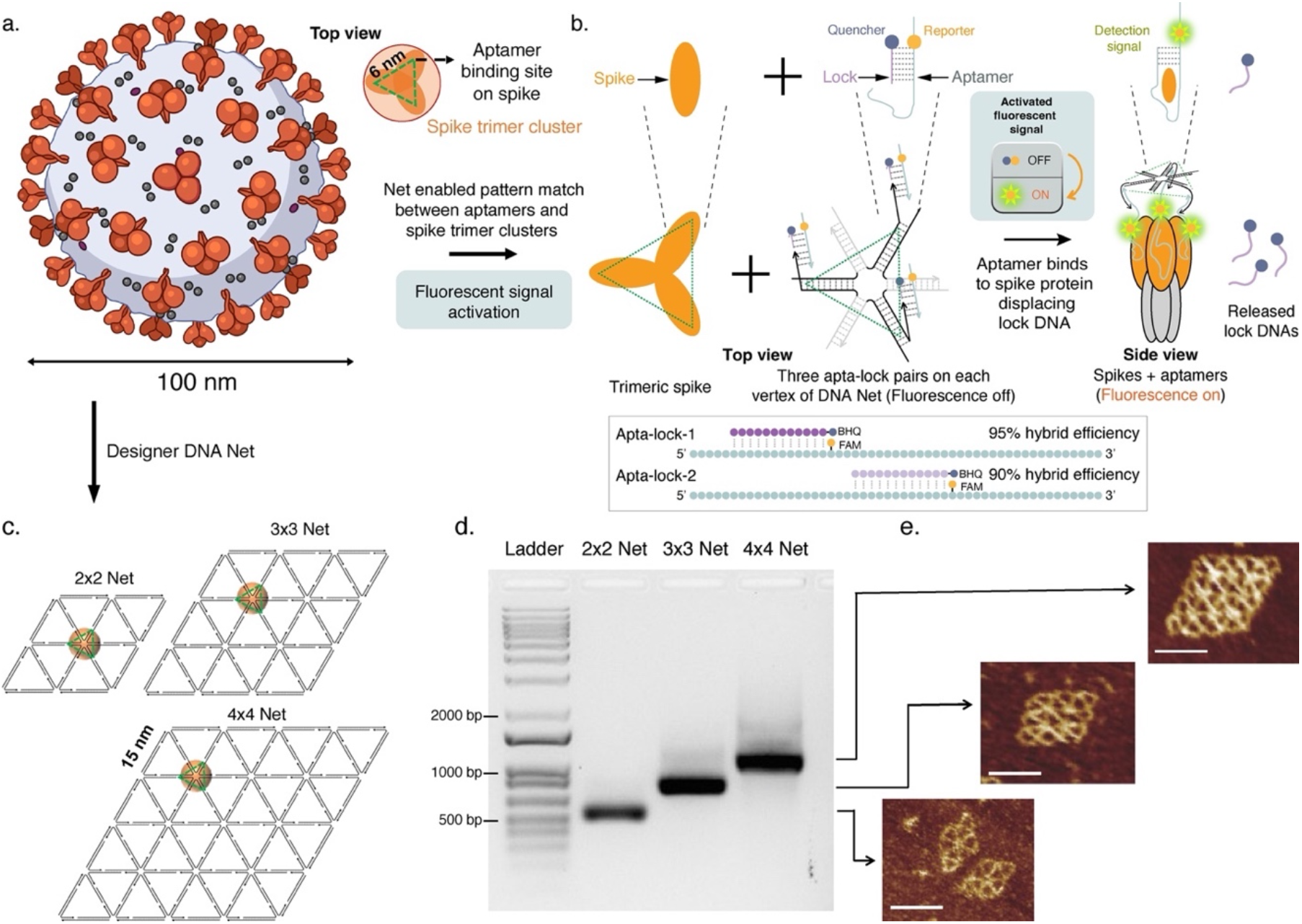
Spike structure, pattern analysis, fluorescent signal activation, and DNA Net design and characterization. **a**, Illustration of SARS-CoV-2 virus. The diameter of a virion is ∼ 100 nm. The spacing between two aptamer-binding sites on the spike RBD within a trimeric cluster is ∼ 6 nm. **b**, Scheme and mechanism of fluorescence signal generation upon SARS-CoV-2 virus interaction with DNA Net-aptamers complex. Top row: schematic of “off-on” aptamer switching at a monomeric interaction level. Middle row: schematic of off-on aptamer switching at trimeric interaction level. Each vertex of the DNA Net consists of three aptamer-lock pairs. The strong aptamer-spike binding triggers the departure of the lock DNA, thus restoring the fluorescence from the fluorophore. Bottom row: Schematic of Apta-lock-1 and Apta-lock-2 designs. **c**, Each DNA Net (2×2, 3×3, and 4×4 Nets) is designed with 3’ end of each DNA oligonucleotide indicated by an arrow. Three aptamer-lock pairs (for sensing) or vanilla aptamers (for inhibition) are placed at each vertex (indicated by a green triangle) to match the intra-spatial pattern of spike trimers. Each orange sphere indicates a trimeric spike cluster. **d**, Formation of the DNA Nets was characterized by 1% agarose gel electrophoresis (AGE). **e**, AFM images of 2×2, 3×3, and 4×4 Nets. Scale bars indicate 30 nm. Supplementary Fig. 1 shows AFM images with a larger scan area.

### Design principle of SARS-CoV-2 virion triggered fluorescence sensing

Nucleic acids have been engineered to act as a controllable switching material in response to external stimuli including ligand-binding, light, pH, and other microenvironmental cues^38^. Herein, we developed a two-strand, ligand-binding triggered aptamer switching motif, inspired by how aptamer switch probe acts^39^, to turn the DNA Net-aptamers into a highly sensitive SARS-CoV-2 biosensor using fluorescence readout. As illustrated in **Fig. 1b**, each aptamer (apta) patterned on the DNA Net is FAM-labeled but its fluorescence is quenched by a BHQ1-tagged “lock” DNA that hybridizes with the aptamer. There are three apta-lock pairs on each vertex of the DNA Net to promote DNA Net-aptamers binding with SARS-CoV-2 surface-displayed spike trimers through pattern-matching (three aptamers with three spikes in each trimeric cluster) and multivalent interactions (multiple tri-aptamers with multiple trimeric spikes on the virus surface custered by the DNA Net scaffold). In presence of spike protein, the binding between the aptamer and the spike will disrupt the aptamer-lock hybridization, resulting in the separation of the quencher from the fluorophore and thus switching on the fluorescence as a virus detection signal. The virus-triggered fluorescence is easily detectable with a portable fluorimeter, as used in this study, or by a bench-top, high throughput microplate reader, such as those used in conventional RT-qPCR instruments, as we previously demonstrated^16^. Equivalent to a LoD concentration in sensing, a minimum spike protein concentration needed to trigger this “off-on” aptamer switching is determined by the aptamer-spike binding affinity. In other words, achieving higher aptamer-spike binding affinities will reduce the spike concentration (equivalent to lowering LoD in sensing) required to reach detectable sensing signals.

We used NUPACK^40^ to design candidate locks that hybridize with the aptamer at different loci based on the predicted secondary structure of the aptamers and the reported nucleotides that interact with spike-RBD residues at the aptamer-spike interface^41^ (**Fig. 1b middle**). Three criteria were taken into the consideration when designing and selecting candidate “apta-lock” pairs for the downstream screening using SARS-CoV-2 sensing: (1) melting temperature of the pair set at a minimum of 45 °C at physiological salt concentrations to ensure that apta-lock duplex won’t fall apart and trigger false positive signals in freshly collected patient samples that could be at 37 °C; (2) Lock DNA does not hybridize with the 5’ or 3’ ends of the aptamer; (3) the apta-lock hybrid efficiency is scored at 90% or higher in NUPACK^40^ so the quencher on the lock can effectively quench the fluorophore on the aptamer by design. In the end, two apta-lock pairs, called Apta-lock-1 and Apta-lock-2, that meet these criteria were shortlisted and further compared and evaluated in the DNA Net-based SARS-CoV-2 sensing test settings (illustrated in **Figs. 1b bottom**).

### Docking and molecular dynamics (MD) simulations of aptamer-spike interactions

We performed docking and MD simulations of the RBD-targeting aptamer with the spike protein to computationally validate our design strategy for SARS-CoV-2 detection. Specifically, we compared the binding modes of free single aptamers versus a tri-aptamer configuration with the spike trimer to decipher if using the latter can increase the -binding affinity and avidity with stronger molecular interactions. Additionally, we also varied the spacing between the constituent aptamers within a tri-configuration to determine the best binding mode. These analyses proved beneficial to our real time assay sensitivity (discussed in the later sections) once the aptamers were arrayed on the DNA Net. Our study shows that a single spike RBD-targeting aptamer binds with a spike protomer in multiple binding modes with binding energies ranging from -10.34 to - 23.9 kJ/mol. The tri-aptamer configuration binds to the spike trimer with binding energies ranging from -68.18 to -72.49 kJ/mol. The highest binding energy configuration for this case has a separation of 6.33 nm between two spike-interacting ends of the constituent aptamers (**Supplementary Fig. 2a**). Additionally, the ends of the aptamers that bind to the DNA Net are spaced at a distance of 5.56 nm among each other (**Supplementary Fig. 2b**). Increasing this distance from 5.56 nm to 9.92 nm reduced the binding energy between the tri-aptamer cluster and the spike protein (**Supplementary Fig. 2c**). RMSD (root mean square deviation) analysis of the MD simulation for the best binding configuration shows negligible variation for the protein residues (**Supplementary Fig. 2d**), suggesting strong and stabilizing interactions between the aptamers and the Spike protomers. Each constituent aptamer exhibits molecular interactions with two Spike protomers and shares a significant portion of the binding surface used by Spike protein to bind to its natural receptor, ACE2R (**Supplementary Fig. 2e**). These interactions include hydrogen bonds, salt bridges and hydrophobic interactions, in addition to multiple Van der Waals interactions (**Supplementary Fig. 2f**).

### Synthesis and characterization of DNA Net nanostructures

We designed different sized DNA Nets containing triangles as the minimal structural/repeating unit and carrying an increasing number of tri-aptamer clusters to achieve an optimum size array for a near RT-PCR sensitivity. We hypothesized an enhancement in the binding strength/affinity of aptamer-spike interactions, assay sensitivity, and inhibition performance for larger size Nets through multivalent interactions enabled by the array of tri-aptamer clusters on each DNA Net. DNA Nets consisting of 2×2, 3×3, and 4×4 individual rhombi were constructed *via* the DNA tile design principle (minimizing DNA sequence symmetry) and assembly strategy^16,19,42^. Each rhombus edge is made of 42 bp DNA (13.6 nm long). The shorter diagonal of each rhombus unit is filled in with a DNA duplex “strut” by forming two triangles to prevent the deformation of the rhombus or in turn the entire DNA Net structure. The triangular repeating units represent an optimal structure to effectively cluster and pack mobile spikes on the viral surface^37^. As suggested by a previous study^27^, 4 thymidine (T) long spacers were used at vertices that connect any two adjacent edges to provide the necessary local flexibility for high yield formation of the desired DNA Net nanostructures. Importantly, each vertex on the Net carries three aptamers (∼ 5.8 nm spacing) aiming to simultaneously bind all three spikes within a trimeric cluster (**Fig. 1c**). Since the length of a 4T spacer is estimated to be ∼1.36 nm long, the center-to-center spacing between two adjacent vertices is estimated to be ∼14.96 nm (13.6 ± 1.36 nm).

The successful formation of the DNA Nets was characterized on 1% agarose gel electrophoresis (AGE) (**Fig. 1d**), and further confirmed by atomic force microscopy (AFM) imaging (**Fig. 1e** and **Supplementary Fig. 1**). The gel also showed presence of DNA Net dimers and multimers in a miniscule amount. However, this should not affect the overall Net’s sensing or inhibition abilities, as the amount is negligible while the multimers still display aptamers in the desired pattern albeit with an enlarged Net size. Notably, unlike DNA origami^43-45^, the thin tile-shaped DNA Net structures can be easily deformed or damaged by deposition on mica, during sample preparation for AFM characterization. We also tested the stability of DNA Nets at reduced temperatures (4 °C or after freeze-thaw cycles) and in saliva matrix to configure and master the conditions for its long-term storage and deployments in clinical applications. Shown by the AGE assays (**Supplementary Fig. 3**), a DNA Net structure that we have intentionally kept with the oldest sample age was proven to be stable after a 1-month (longest we tested) storage at 4 °C, and after at least three freeze-thaw cycles between -20 °C and the room temperature in the laboratory (23 °C). The DNA Net also shows no degradation following 1 h (longest we tested) of incubation in saliva at room temperature (**Supplementary Fig. 4**).

Next, we performed surface plasmon resonance (SPR) study to confirm that the DNA Net aptamers dramatically improved the binding affinities compare to solitary aptamers, and larger DNA Net size can result in higher binding affinities through increased aptamer-spike binding events. **Supplementary Fig. 5** shows that the binding strength of the aptamer to the spikes increased by 1 × 10^3^-fold when the same aptamers were patterned on 2×2 DNA Net, or by 1 × 10^6^-fold on 4×4 DNA Net. We also observed that DNA Nets with (DNA Net Apta-lock) or without the lock DNA (DNA Net Aptamer) exhibited similar binding affinities to the trimeric spike proteins on the SPR chip, confirming that DNA lock, as part of the signal reporting machinery in our DNA Net sensor, does not compromise the binding between spike and the DNA Net-aptamers (**Supplementary Fig. 6**). Thus, the resulting improvement in the binding avidity, in turn, led to high sensitivity using our DNA Net sensor, as validated in our subsequent assays.

### Enabling multivalent aptamer-spike protein interactions enhances SARS-CoV-2 detection sensitivity

Our previously reported study on Dengue virus detection using ‘DNA star’ sensor demonstrated that higher degrees or valency of “global” pattern matching between DDN-arranged aptamers and antigen clusters on the virus will lead to stronger ligand-antigen binding affinities. This resulted in higher detection sensitivity as well as inhibition potency for our sensor^16,19^. Our SPR assay also confirmed that increasing the number of aptamer-spike binding sites through an increase in the DNA Net size results in a stronger binding affinity to the trimeric spikes on SPR chip. Thus, for our fluorescence-based DNA Net assay, we first selected the best aptamer-lock candidate between the Apta-lock-1 and Apta-lock-2 using 4×4 DNA Net (**Fig. 2a**). Our sensing assays showed that DNA Net sensor with Apta-lock-1 generated a significantly better signal intensity and sensitivity than Apta-lock-2 (**Fig. 2b**). As a result, the ensuing studies reported henceforth were conducted using Apta-lock-1 as the reporter unit on the DNA Net sensor for SARS-CoV-2 detection.

**Fig. 2.**
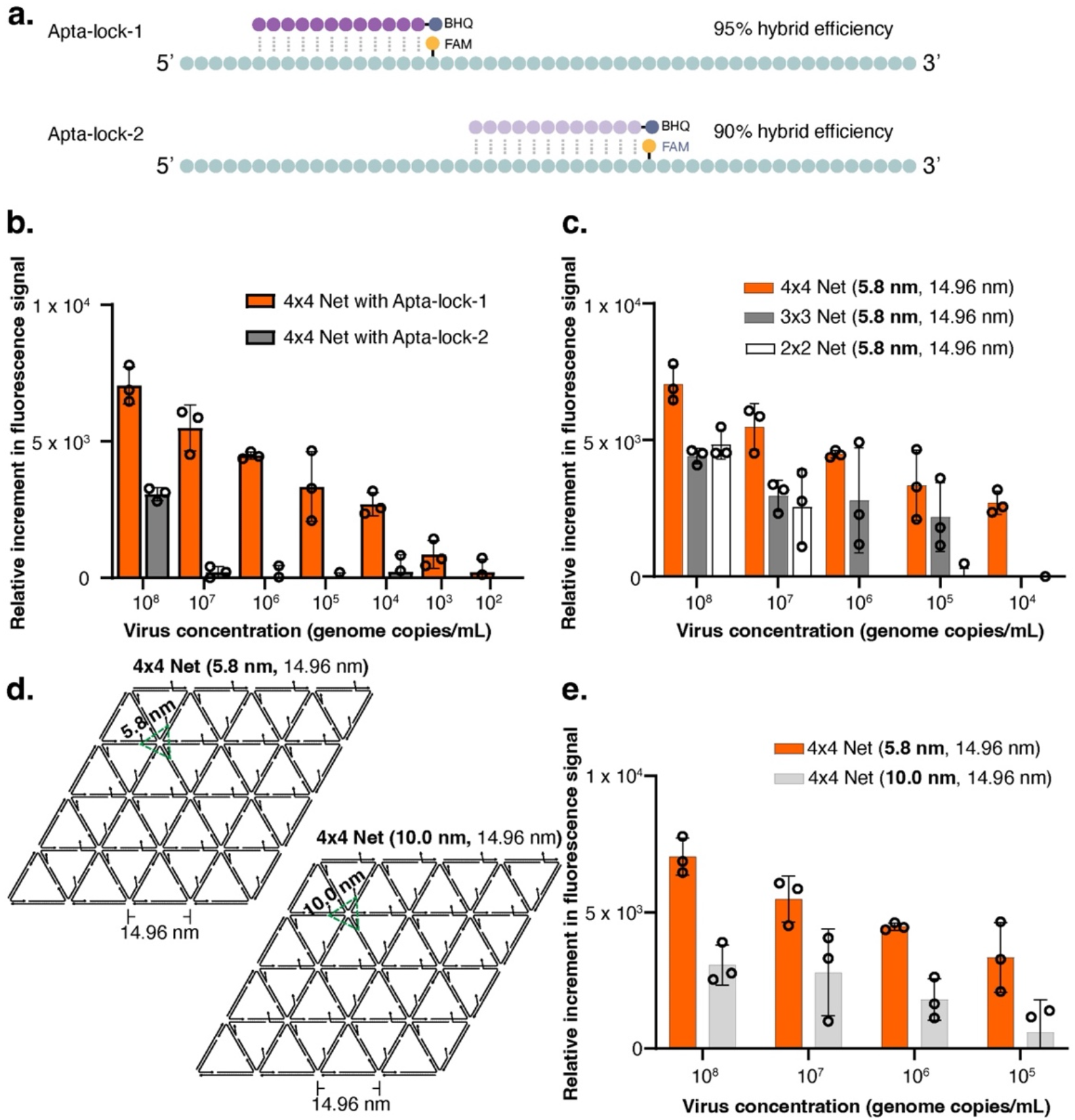
Design and characterization of aptamer-lock pair, Net size, and impact of intra-tri-aptamer spacing on sensitivity of DNA Net sensors. **a** Schematic of Apta-lock-1 and Apta-lock-2 designs. **b**, Detection sensitivity of 4×4 DNA Net sensor with Apta-lock-1 (LoD is 1×10^3^ viral genome copies/mL) or Apta-lock-2 (LoD is 1×10^8^ viral genome copies/mL) with pseudotyped SARS-CoV-2 virus. **c**, Detection sensitivity of 2×2 Net, 3×3 Net, and 4×4 Net sensors with pseudotyped SARS-CoV-2 virus. **d**, Schematic of DNA Nets with designed (intra-, inter-) tri-aptamer spacing. **e**, Detection sensitivity of 4×4 DNA Net (**5.8 nm**, 14.96 nm) and 4×4 DNA Net (**10 nm**, 14.96 nm) with pseudotyped SARS-CoV-2 virus. In **b, c, e**, data are presented as mean ± s.d., n = 3 biologically independent samples. Individual data points below background are not shown but were involved in error calculation.

Furthermore, we tested the performance of the 4×4 DNA Net sensor in two solution matrices, in 1 x TA-Mg^2+^ buffer (sensing buffer), or in artificial saliva^46^. The former could be a liquid medium used to elute virus samples collected on nasopharyngeal swabs for onsite detection. Our DNA Net sensor performed equally well in both the sensing buffer and saliva matrices that contain high viral loads (**Supplementary Fig. 7**). At low viral loads, the DNA Net sensor works marginally better in artificial saliva, possibly because SARS-CoV-2 virus are more compatible in a medium that mimics saliva. Thus, we characterized the DNA Net sensor in artificial saliva containing solution for the remainder of the study.

To determine the effect of increasing multivalent interactions in enhancing the detection sensitivity of DNA Net sensor for SARS-CoV-2, we systematically performed the detection assays starting from the monomeric Apta-lock pair, 6-arm junction structure (representing a single tri-aptamer cluster on a vertex of DNA Net), and different size DNA Nets (2×2, 3×3, and 4×4) that each carries an increasing number of apta-lock pairs after normalizing the apta-lock concentration. The monomeric Apta-lock pair, 6-arm junction structure (**Supplementary Fig. 8**), and the 2×2 DNA Net sensor can detect SARS-CoV-2 but only at extremely higher viral loads, while the 3×3 and 4×4 Net sensors were able to detect SARS-CoV-2 virions with concentration-dependent signal intensities and sensitivities at normalized aptamer concentration (**Fig. 2c**). The detection signal from the DNA Net sensor was obtained within 10 mins following a simple sample “mix-and-read” procedure with a portable fluorimeter (AND1100^28^). The limit of detection reported for most of the current FDA-approved RT-PCR tests is between 1×10^2^ – 1×10^5^ viral genome copies/mL (e.g., Ref^47,48^). Remarkably, the sensitivity of the 4×4 DNA Net sensor (1×10^3^ copies/mL) meets the requirement for high sensitivity in identifying patients at early stages of infection^47,48^.

Next, we tested the concentration dependent sensing performance of monomeric Apta-lock and 6-arm junction sensors to improve the detection sensitivity for these structures. Of note, only these two sensors were used for this purpose because increasing the concentration of all DNA Nets beyond a certain limit resulted in precipitation which is detrimental to the fluorescence read out. Our assay, results of which are presented in **Supplementary Fig. 8**, showed that for both sensors the virus detection sensitivity has a negative correlation with the sensor concentration. At higher sensor concentrations, we observed worse performances of these structures. The assays also demonstrated that within the concentration range tested, both, monomeric apta-lock and 6-arm sensors, display lower detection sensitivities than 3×3 and 4×4 DNA Net sensors.

Further, we investigated whether a larger DNA Net than the 4×4 Net can further improve the detection sensitivity. For this purpose, we synthesized and tested a 5×5 DNA Net sensor with 5.8 nm intra-trimeric aptamer spacing and 14.96 nm unit size, as those of 2×2 to 4×4 Nets. We observed a 10-fold reduction in detection sensitivity for this net size (**Supplementary Fig. 9**). This loss of sensitivity can be attributed to two reasons. First, a poor signal to noise ratio was observed for the 5×5 Net sensor due to a high background signal from the greatly increased number of FAM-tagged aptamers affecting the resolution of signal at low virus concentrations. Second, difficulty to maintain a perfect stoichiometry among the 170+ DNA oligos that form the 5×5 DNA Net contributes to more non-effective sensor molecules in comparison to the 4×4 DNA Net sensor. This also agrees with the decreased sensitivity observed for the Apta-lock and 6-arm junction sensors at higher concentrations. Separately, we also designed and synthesized an overall larger 4×4 DNA Net that carries a larger inter-aptamer cluster spacing (52 bp DNA) but still maintained the 5.8 nm intra-trimeric aptamer spacing (**Supplementary Fig. 10a-b** for design, and AGE/AFM characterization). The center-to-center spacing between two adjacent vertices of the 52 bp unit DNA Net is estimated to be ∼18.36 nm. For 52 bp unit size DNA Net sensor (5.8 nm, 18.36 nm), we observed a slight reduction in the detection sensitivity than 42 bp unit DNA Net (5.8 nm, 14.96 nm) (**Supplementary Fig. 10c**). A possible explanation for the reduction in the detection sensitivity could be that the 42 bp unit 4×4 DNA Net offers the smallest size DNA nanostructure that can maximize the clustering efficiency of the mobile spike proteins as well as provides lesser steric hindrance that results in a higher binding efficiency and sensitivity as compared to the 52 bp unit size 4×4 DNA Net.

To compare our assay sensitivity with gold standard antigen test methods, we carried out an ELISA assay using a commercially available kit to detect purified SARS-CoV-2 spike proteins. The ELISA assay showed an LoD of 15 – 31 pg/mL trimeric spike proteins, which is estimated equivalent to ∼ 6.25 × 10^5^ – ∼ 1.29 × 10^6^ viral genome copies/mL, assuming no loss upon spike protein cleavage and recovery from SARS-CoV-2 virus samples (**Supplementary Fig. 11**). The purified spikes afforded higher sensitivity in our ELISA assay as the sample preprocessing, skipped in our ELISA assay, is considered a major step contributing to the loss of viral antigens reducing the actual sensitivity of the assay.

To summarize, the LoD of our 4×4 DNA Net sensor is comparable to clinically relevant SARS-CoV-2 viral loads (1×10^4^-1×10^10^ viral genome copies/mL in the upper respiratory tract) in patients with or without symptoms^49-51^. Thus, being able to detect a patient’s viral load at an early infection stage is highly beneficial to individual patients and for timely screening and control of pandemic outbreaks within surveillance and diagnostic networks.

### Intra-trimeric spike pattern matching by DNA Net-aptamers complex is important for achieving high SARS-CoV-2 detection sensitivity

Our DNA Net (5.8 nm, 14.96 nm) sensor design utilizes a pattern matching strategy by targeting intra-(6 nm) spike trimeric cluster spatial patterns and effectively clustering spike trimers aiming for maximum binding avidity and detection sensitivity. Herein, we further tested the performance of a DNA Net that has the same inter-trimeric aptamer spacing (42 bp or 14.96 nm) but with an enlarged, 10.0 nm intra-trimeric aptamer root spacing (**Fig. 2d, Supplementary Fig. 12** for the DNA Net design at a single vertex). We observed a decrease in detection sensitivity for this 4×4 DNA Net (10 nm, 14.96 nm) that has an enlarged intra-aptamer spacing, correlating with the decreased binding energy predicted in our in-silico analysis (**Fig. 2e**). However, it still provides a reasonably good sensitivity falling in the clinically relevant range for viral loads, as predicted by our in-silico computational studies (**Supplementary Fig. 2c**), in which the three aptamers arranged at each Net vertex can flex inwards allowing some of which to still simultaneously interact with the 6 nm spaced spikes in the trimeric cluster. Taken together, our DNA Net design (with 5.8 nm, 14.96 nm spacing) principle/strategy was found to be optimal for achieving high detection sensitivity.

### Higher DNA scaffold pliability increases multivalent aptamer-spike interactions to improve virus detection sensitivity

Carrying hollow structures, DNA Nets shall have high mechanical pliability to form a concave shape in solution, thus promoting the multivalent and pattern-matching interactions between the patterned aptamers and the spike proteins on spherical SARS-CoV-2 virions. To further investigate the impact of sensor scaffold bendability on virus detection sensitivity, we designed and synthesized a sensor based on the DNA origami plate^52^ template, which carries a non-hollow structure, and compared its structural bendability and sensing performance with the 4×4 Net sensor. Specifically, we used oxDNA to implement coarse-grained simulations to compare the bendability of two scaffolds as measured by Root mean square fluctuations (RMSF). Shown in **Supplementary Figs. 13 and 14a**, and **Supplementary Movies 1-2**, 4×4 Net displays more wrinkles, or has larger RMSF, than the DNA origami plate. Additionally, the dsDNA edges of the 4×4 Net exhibit larger displacement, while the DNA origami plate tends to keep a planar shape during the entire simulation time. These simulation data indicate that the 4×4 DNA Net scaffold has higher pliability than the DNA origami plate scaffold. Performance comparison of the DNA origami plate sensor (**Supplementary Fig. 14b**) provides a ∼1×10^3^-fold higher LoD (**Supplementary Fig. 14c**) than the 4×4 DNA Net sensor. As expected, higher mechanical pliability of the DNA Net scaffold can facilitate the formation of concave shapes in solution, thus promoting and maximizing the multivalent and pattern-matching interactions between the DNA Net-patterned aptamers and the spike proteins on spherical SARS-CoV-2 virus particles. However, the DNA origami plate despite holding the exact number of aptamers, the rigid (less bendable) nature of the scaffold prevents more individual aptamer units at the plate-virus interface from interacting with multiple spikes on the virus surface.

### DNA Net sensor specificity studies

To explore if the spike-binding aptamer detects the WT SARS-CoV-2 spike epitopes just because of its physical characteristics (like size, orientation, etc.) match or by a chemical or molecular match, we performed specificity tests for the DNA Net sensor against H1N1 (influenza) and OC43 (a common cold human coronavirus). These viruses carry class I fusion proteins in trimeric clusters, like the spike trimers of SARS-CoV-2 virus but with different surface antigen spacing, with different antigen amino acid composition for our aptamer to target. This was also important for the selectivity of DNA Net sensor in clinically relevant settings since both influenza A (H1N1) and human coronavirus OC43 are common respiratory viruses that present similar respiratory symptoms as COVID-19. We also tested the sensor specificity against Zika virus, as an additional control. If the DNA Net sensor does not recognize a virus particle through proper interaction mediated *via* matched pattern or binding aptamer, the fluorescent reporters will remain quenched. Our results verified that the 4×4 DNA Net sensor designed for SARS-CoV-2 detection does not cross-react with H1N1, OC43, or Zika, even at high viral loads (**Fig. 3a**). These data suggest that our SARS-CoV-2 biosensor is selective and that our DNA Net-based virus sensing strategy can be adapted for multiplexed detection of commonly circulating respiratory viruses by using specific aptamer-lock pairs and fluorophores with easily distinguished emission wavelengths for signal readout.

**Fig. 3.**
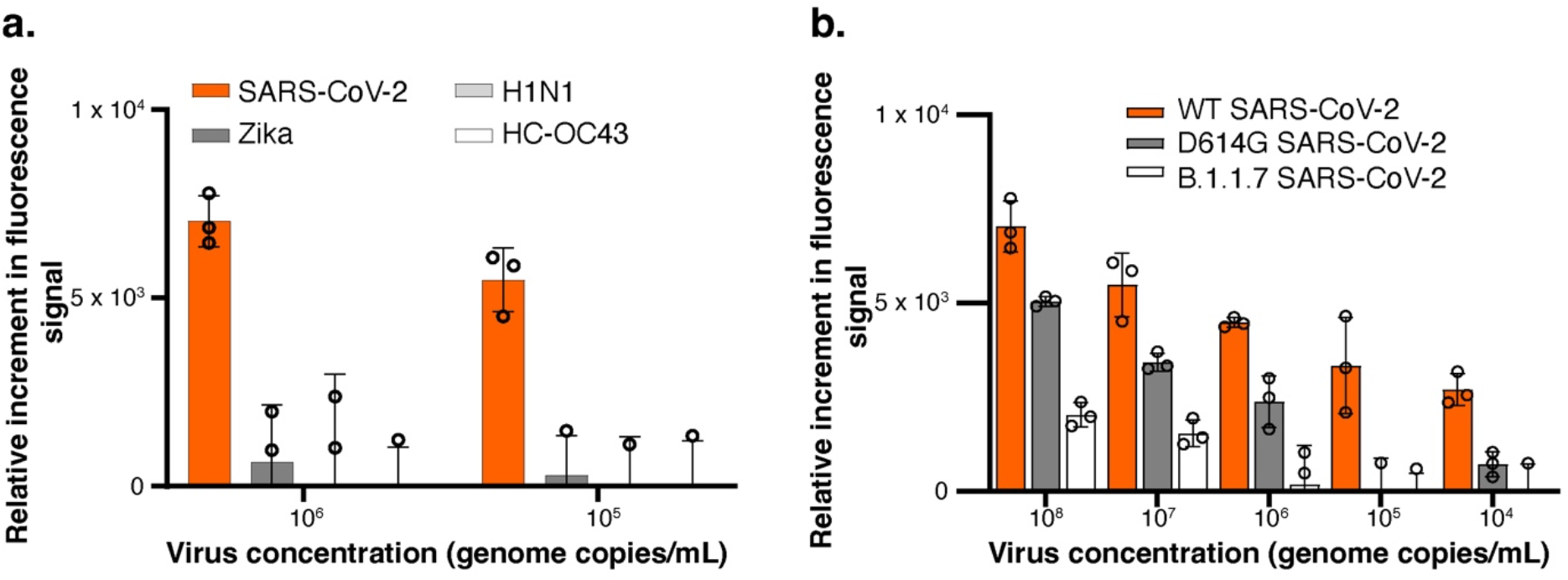
Selectivity of DNA Net sensor. **D**ata are presented as mean ± s.d., n = 3 biologically independent samples. Individual data points below background are not shown but were involved in error calculation. **a**, Cross reactivity study for the 4×4 DNA Net sensor with H1N1 (an influenza-A virus), Zika virus, and OC43 (a human coronavirus). **b**, Detection sensitivity of the 4×4 DNA Net with D614G and B.1.1.7 SARS-CoV-2 mutants.

### Detection of SARS-CoV-2 variants by DNA Net sensor

SARS-CoV-2 is constantly evolving, with emerging variants that are highly transmissible and gradually becoming dominant in circulation. To further determine whether our DNA Net sensor which carries an aptamer selected against WT SARS-CoV-2 spike can detect the emerging SARS-CoV-2 mutant strains in human population at the time of this study, we tested the 4×4 DNA Net in detecting pseudotyped viruses that resemble the D614G or B.1.1.7 mutants. We found that our DNA Net sensor can detect both variants but with lower signal intensities and sensing sensitivities (**Fig. 3b**). The LoD was ∼ 1×10^6^ viral genome copies/mL for detecting D614G, and ∼ 1×10^7^ viral genome copies/mL for detecting B.1.1.7. Both of the D614G^53^ and B.1.1.7 mutant^54^ viruses exhibit elevated replication and/or shedding in infected individuals, compared to the WT strain. The D614G mutation on the spike has been reported to alter the spike trimer interactions^55^ resulting from altered spike protein spatial patterns, which may affect the sensitivity of our DNA Net sensor. For the B.1.1.7 lineage, its spike protein contains 69/70 deletion that likely leads to an even more conformational changes in the spike architecture^56^, leading to a reduced Net detection sensitivity.

### DNA Net can distinguish infectious SARS-CoV-2 from its inactivated counterparts

The performance of our DNA Net sensor was first screened and demonstrated using pseudotyped models of the WT and mutant SARS-CoV-2 strains. These pseudotyped viruses reconstitute the spike protein features of active SARS-CoV-2 virus, are readily accessible to researchers, and can be handled safely in a laboratory with biosafety level 2 (BSL 2) containment, which makes these ideal for wide use in the COVID-19 research community ^57-59^. Subsequently, the real, authentic SARS-CoV-2 virions were then used to confirm a similar DNA Net sensing performance by following the same assay procedures as used in detecting pseudotyped viruses, and to show that DNA Net can distinguish infectious SARS-CoV-2 from its inactivated counterparts. Shown in **Fig. 4a**, the best-performing 4×4 Net sensor identified above can detect the authentic live SARS-CoV-2 with a LoD of 1,000 viral genome copies/mL, same as that obtained when using pseudotyped viruses. Since our DNA Net is targeting spike proteins on intact virus particles for the best sensing performance, it should be able to distinguish infectious from heat or chemically inactivated viruses, which possess denatured and cleaved spike proteins and thus are not infectious anymore. Our assays showed that the DNA Net sensor cannot sensitively detect heat or chemically inactivated authentic virus at very high viral concentrations (**Figs. 4b**).

**Fig. 4.**
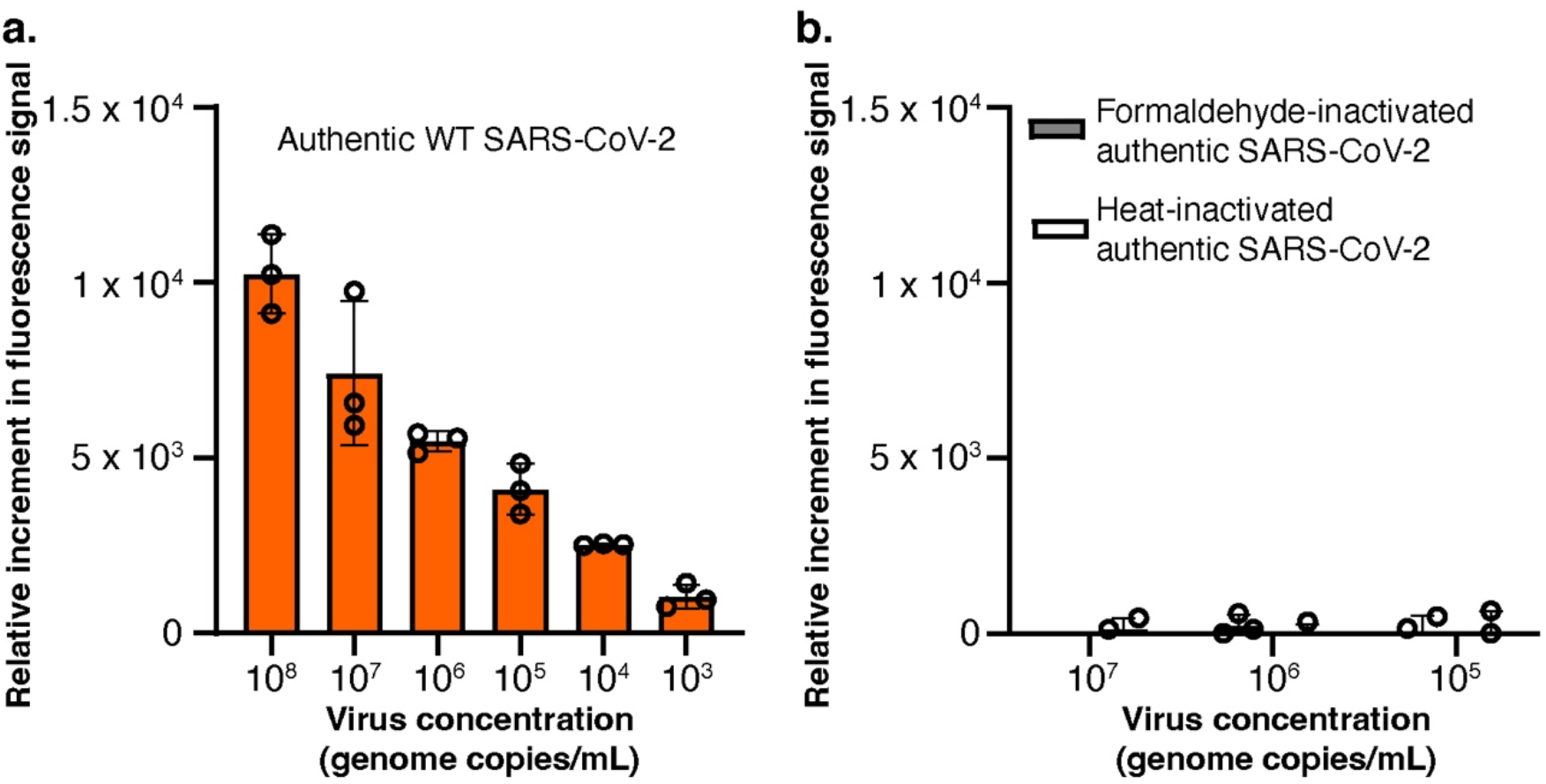
DNA Net sensing of authentic SARS-CoV-2 viruses. Data are presented as mean ± s.d., n = 3 biologically independent samples. Individual data points below background are not shown but were involved in error calculation. **a**, Detection sensitivity of DNA 4×4 Net sensor with authentic SARS-CoV-2 virus. **b**, Detection sensitivity of DNA 4×4 Net sensor with heat-inactivated authentic SARS-CoV-2 virus or chemically inactivated authentic SARS-CoV-2 virus.

### Time-lapsed confocal imaging

The portable fluorimeter used to measure the fluorescence signal from the 4×4 DNA Net sensor reads an ensemble signal in bulk solution. Studying the DNA Net sensor interaction with the virus at a single molecule level in solution typically requires a sophisticated imaging technology such cryo-EM which is challenging given instrument limitation, low contrast signal of thin 2D DNA Net nanostructure, and synthetic limit to the pseudotyped virus concentration. As an alternative, we sought to utilize confocal microscopy to visualize the detection of SARS-CoV-2 virus at a single Net sensor level.

Specifically, for the DNA Net mediated surface capture assay the coverslip surface was functionalized with 0.2 µM amino labeled-DNA Net sensors extended from a corner of the Net. On this functionalized coverslip, a PDMS gasket with two circular openings was placed to form two independent liquid compartments namely Well-1 and Well-2. Well-1 (shown in green in **Fig. 5a**) served as a testing region for active SARS-CoV-2 sample, while heat-inactivated SARS-CoV-2 virus was added into Well-2 (shown in blue in **Fig. 5a**) for “infectious vs. noninfectious” selectivity test. The confocal images were taken at the bottom of the coverslip where DNA Net sensors were immobilized with an 100X oil-immersion objective lens prior to and after addition of SARS-CoV-2 sample.

**Fig. 5.**
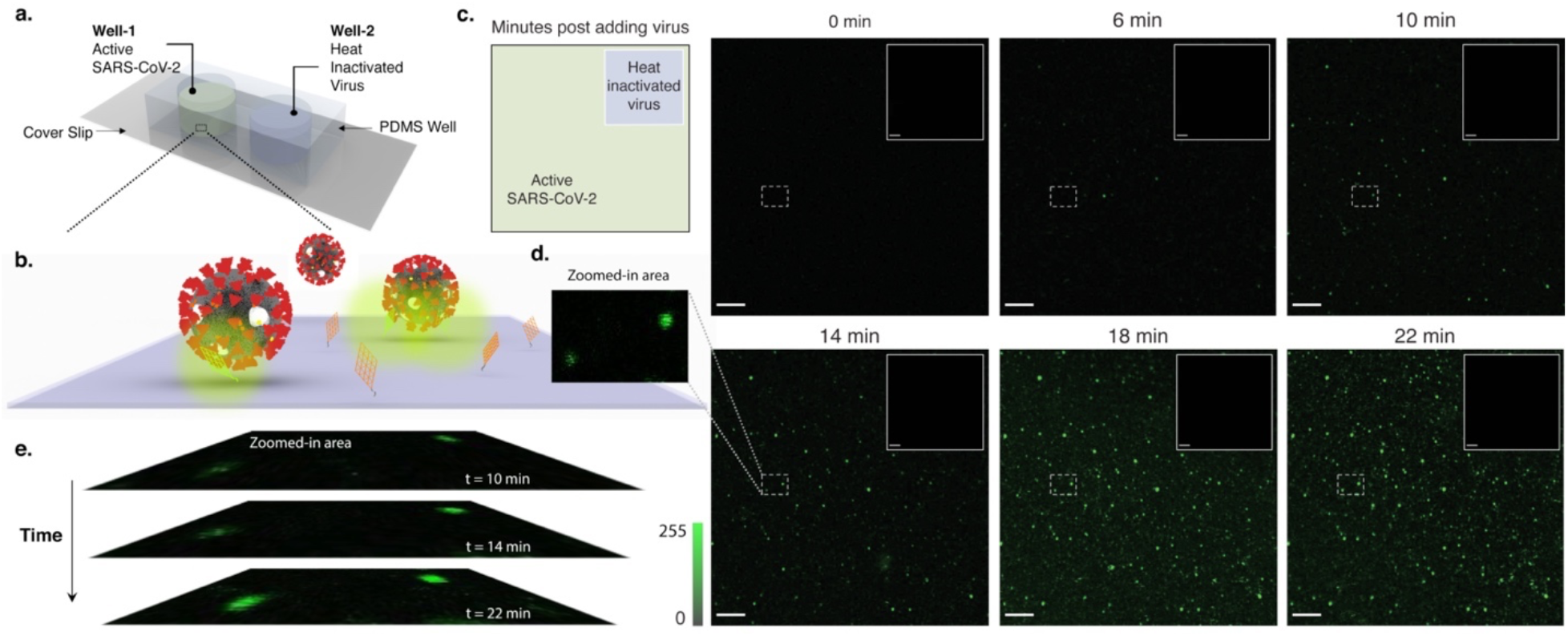
Confocal imaging for SARS-CoV-2 virus detection at single Net sensor level. **a**, Schematic illustration of surface-based sensing assay in two separate testing regions (PDMS wells). **b**, Schematic illustration of DNA Net sensor mediated glass surface capture of active SARS-CoV-2 virions. **c**, Line-scanning fluorescent confocal images for surface-attached DNA Net sensor (4×4) reacting with SARS-CoV-2 virions or heat-inactive SARS-CoV-2 viruses (inset). Scale bar indicates 10 mm. Laser excitation: 488 nm. Emission filter: 505-560 nm. The Zeiss LSM 710 confocal microscope with Argon 488 nm laser, pinhole size: 68 mm. **d**, Zoomed-in area of the image contains two diffraction limited fluorescent spots. **e**, Reaction kinetic response at different timepoints.

Under the confocal microscope, each 4×4 DNA Net sensor results in diffraction-limited bright spots when the quencher-labeled lock DNAs are released during the binding events between the aptamers patterned on the DNA Net and the spikes on the outer surface of SARS-CoV-2 virions. Illustrated by **Fig. 5b**, one SARS-CoV-2 virion could be captured by at least one DNA Net sensor on the surface. The time-lapsed confocal images showed that the fluorescence signal is detected 6 mins after the SARS-CoV-2 sample was added (**Fig. 5c)**. Subsequently, the number of diffraction-limited bright spots increased over time since more viruses could diffuse to the surface to interact with the DNA Net sensors. At the same time, the fluorescent intensity of each diffraction-limited spot also increased progressively in the course of time (**Fig. 5e**). This was expected, as with time, the number of spike-aptamer interactions between the DNA Net and virus increases releasing vast quantities of quencher DNA at the approximate location within diffraction limits. Conversely, there were no detectable fluorescent signal changes under the scanning region in well-2 after 22 mins (the longest time we tested as a comparison) following addition of the heat-inactivated SARS-CoV-2 sample. Our time-lapsed confocal assays provide a direct imaging evidence that the DNA Net sensor can detect the presence of SARS-CoV-2 and distinguish between the infectious and the noninfectious forms of the virus. However, monovalent Apta-lock immobilized on the surface using the same preparation (illustrated in **Supplementary Fig. 15**) failed to generate any fluorescence signal even after 24 h of incubation with high concentration of SARS-CoV-2 virus. Of note, it is expected that the rate of Net-virus interaction on surface is lower than in free solution since the substrate surface can cause steric hindrance to block and decelerate the Net-virus interactions.

### DNA Net in SARS-CoV-2 inhibition

In our previous study, the “DNA Star” could not only bind to DENV with high selectivity and specificity, but also effectively block DENV infection^16,19^. In our DNA Net-aptamers assay, the monomeric binding unit is the spike RBD-binding aptamer, which was organized in a trimeric fashion to mirror the intra-spike trimer spacing pattern, and tri-aptamers were further organized on the DNA Net to achieve maximum aptamer-spike interaction sites. As suggested by our in-silico analysis and validated by our SPR assay, the elevated avidity of the DNA Net-aptamers compared to free aptamers (monomeric control) contribute to an enhanced binding and can potentially exhibit inhibition capability against SARS-CoV-2 spike-mediated infection. To determine whether our DNA Net-aptamers designed for detecting enveloped viruses could also block viral infection, we performed an inhibition assay using the authentic live SARS-CoV-2 infected VERO-E6 cells and quantitated the antiviral efficacy of the DNA Net-aptamers by using high throughput flow cytometry detecting the intracellular SARS-CoV-2 nucleocapsid protein (**Fig. 6a**). Interestingly, we found that the aptamer can impede SARS-CoV-2 infection with an IC_50_ ∼ 10.70 µM, and strikingly, the 4×4 DNA Net-aptamers has an IC_50_ at of approximately 11.81 nM (95% confidence interval between 7.32 nM to 20.26 nM) (**Fig. 6b**), representing nearly 1×10^3^-fold enhancement in anti-viral inhibition compared to the monomeric aptamer. DNA Net-aptamers at the IC_50_ concentration did not exhibit any cytotoxicity compared to the cell medium (**Supplementary Fig. 16**). Anti-SARS-CoV-2 efficacy increases as the number of aptamers per DNA Net scaffold increases from 2×2, 3×3 to 4×4(**Supplementary Fig. 17**), while the DNA Net alone does not exhibit any antiviral activity (**Fig. 6b**). Thus, in additional to being an excellent detection platform, the performance of the DNA Net-aptamers against *in vitro* SARS-CoV-2 inhibition shows a promising therapeutic potential.

**Fig. 6.**
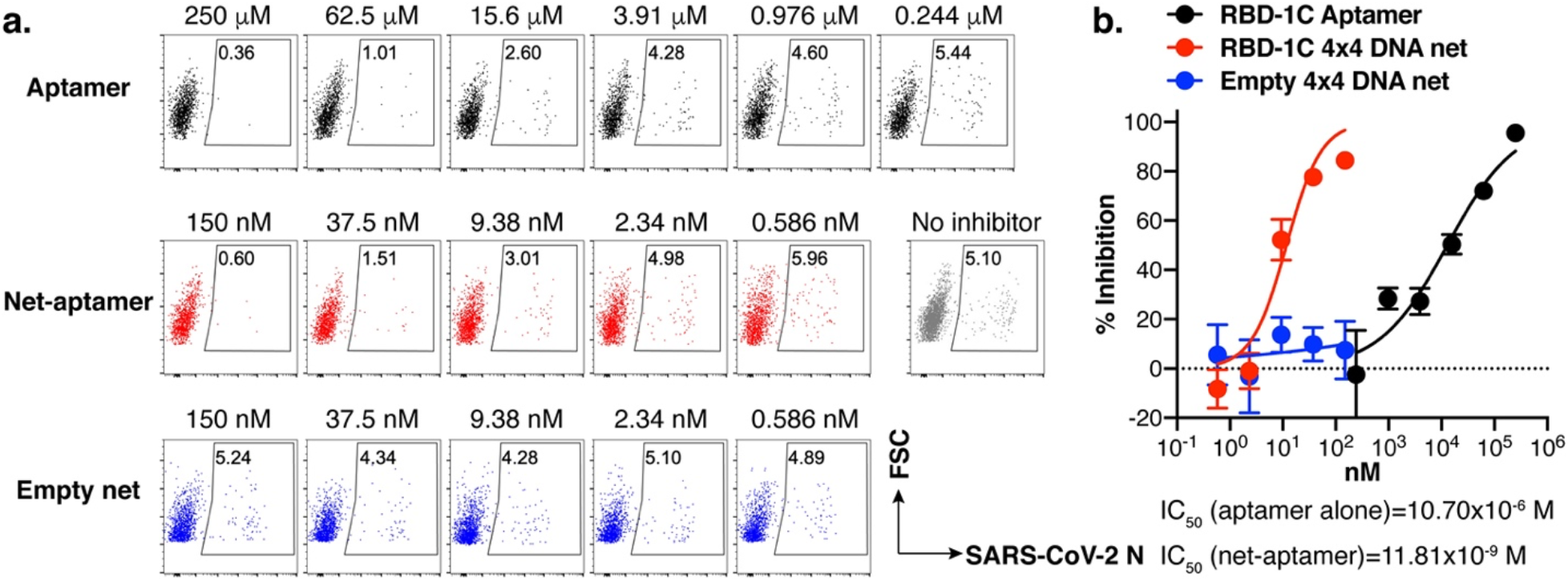
Inhibition of active SARS-CoV-2 infection in VERO-E6 cells. **a**, Representative flow cytometric plots of intracellular SARS-CoV-2 N protein staining in VERO-E6 cells infected with USA-WA1 (MOI = 0.05). **b**, Nonlinear regression was performed using the inhibition curves to calculate the IC_50_ and the 95% confidence interval of the aptamer (IC50: 10.70 µM; 95% CI: 6.56 µM to 16.98 µM), 4×4 DNA Net-aptamers (IC50: 11.81 nM; 95% CI: 7.32 nM to 20.26 nM), and 4×4 DNA Net without aptamer (IC50: N/A, 95% CI: N/A). Data represent results of two independent experiments and are presented as mean ± s.e.m., *n* = 3 biologically independent samples.

## DISCUSSIONS

The challenges for COVID-19 diagnosis are typical and historically evidenced by repeated tragedies during previous emerging epidemic and pandemic infections, especially in detecting etiologic RNA viruses for most of the high-impact human viral diseases^60,61^. Diagnosing infectious diseases caused by emerging RNA viruses, reduced sensitivity of PCR-based tests can result from the low amount of starting material (1 RNA genome copy per SARS-CoV-2 viral particle), loss of RNA during the extraction process, and instability of the extracted RNA. Our DNA Net sensor shows superiority over gold standard antigen tests. It exhibited a range of detection sensitivity among those of the PCR-based tests. This was achieved through the selectivity of spike-targeting aptamer and in rational design of DNA Net to allow matching intra-trimeric aptamer spacing to the spike proteins on the viral surface and to enable multivalent spike-aptamer interactions. In principle, multiple DNA Nets can bind to a single SARS-CoV-2 virus particle for the simultaneous release of many fluorescent reporters from a single virus particle binding event (**Scheme 1**) resulting in detectable signal readout even at low viral concentrations. This gives the DNA Net platform an edge over the current and existing aptamer-based detection platforms that depend on monovalent or non-pattern matching interactions which results in lower sensitivity of such methods. The 4×4 DNA Net sensor with 42 bp unit size (14.96 nm for maximum spike clustering) and 5.8 nm intra-trimeric aptamer spacing (matching to SARS-CoV-2 intra-trimeric spike spacing) gave us the best sensitivity with a LoD of 1,000 viral genome copies/mL in the saliva containing solution, and showed no cross-reactivity against the other common respiratory viruses. The range of detection sensitivity of our DNA Net sensor is among those of the PCR-based tests, while the DNA Net sensor has direct access to the intact viruses in the sample. Thus, our approach circumvents the need to extract nucleic acid material from the virus, RNA purification, enzymatic amplification, thermal cycles, and calibration of the complex equipment resulting in a quick sensitive detection assay that can function isothermally at room temperature and provides an easily interpretable result in the POC settings. Cost of the DNA Net sensor is ∼ $1.26 per test (**Supplementary Table 1** for the cost calculation) which is much lower than ELISA based antigen tests in laboratory settings (∼ $5-10 per test) or lateral flow assay (LFA) based antigen tests (∼ $20 per test)^62^. Additionally, our approach, by directly sensing intact SARS-CoV-2 virions, may also address a problem for people being able to know when they are no longer infectious and can come out of quarantine, as nucleic acid tests are known to generate false positive results from the presence of nucleic acid molecules from degraded viruses.

Additional to detection, the DNA Net-aptamers exhibited promising therapeutic potential during *in vitro* SARS-CoV-2 inhibition. The theoretical free energy of binding between the trimeric spike (−72 kJ/mol) and the tri-aptamer configuration calculated in our in-silico studies was found to be higher than the experimentally determined free energy of binding between the trimeric spike and its natural cognate receptor, the ACE2 (−59 kJ/mol), which explains the inhibitory effect of the aptamer on viral infection. It is known that many viruses first interact with negatively charged glycosaminoglycans (GAGs) on the host cell surface before invasion^63-67^. Thus, the DNA Net-aptamers not only relies on specific polyvalent interactions for binding SARS-CoV-2 but also electrostatically traps and isolates virions from the host cell plasma membrane and GAGs through the negative charges of the DNA scaffold. Additionally, our DNA Net-based strategy can be used to deploy aptamers into arrangements that increase specific, on-target binding while reducing off-target binding based on pattern identity and/or epitope clustering. We achieved a ∼ 906-fold antiviral efficacy enhancement of the same monomeric aptamer using our 4×4 DNA Net-aptamers assay. A similar conclusion was reported using spherical neutralizing aptamer-gold nanoparticles (SNAP) conjugates that resulted in a ∼ 356-fold IC_50_ enhancement of the same monomeric CoV2-1C aptamer by attaching to the AuNPs to form CoV2-1C-AuNP complex^68^. This further echoes with our claim on the importance of pattern matching and multivalent interactions for achieving high antiviral efficacy, elucidating the potential of our platform for viral inhibition and its better antiviral performance over the current state-of-the-art techniques.

Our DNA Net could potentially complement the function of neutralizing antibodies (Nabs, normally with a positive charge) and capture Nabs-escaped viruses by still providing Nabs access to viral epitopes otherwise electrostatically shielded from host cells. Interestingly, most class I and class II human Nabs against the spike protein exhibit binding free energies ranging from 60-80 kJ/mol (ref.^69^), highlighting the selectivity, specificity, and affinity of the DNA Net-aptamers binding for spike protein, and further hints at its potential as a possible therapeutic agent. The SPR and antiviral assays show the DNA Net offering pattern-matching and multivalent interaction can greatly improve the binding affinity and in turn the antiviral efficacy of a viral surface antigen binder (e.g., a spike-binding aptamer used in this study), despite the monovalent binder being a relatively weak binder and a lower potency inhibitor. We speculate that our DNA Net can provide synergy by also turning an already strong binder into an even stronger binder for further improved performance in viral inhibition, as demonstrated by previous studies, in which multivalence with average spacing-matching, or with spatial pattern-matching is utilized to turn a weak (e.g., oligosaccharides, ref.^63^) or an already strong (e.g., nanobody, ref. ^70^) binder into a stronger binder. Of note, although DNA nanostructures have shown stability *in vivo*^71,72^, they can be UV crosslinked^73^ and/or coated with biocompatible ligands (i.e., PEGylated lipids^74^, bovine serum albumin^75^, PEGylated oligolysines^76,77^) to further improve its *in vivo* stability by reducing risks of nuclease degradation and/or low salt denaturation^78-80^.

To summarize, we have created a versatile, sensitive, single step “direct” viral recognition platform based on Net-shaped designer DNA nanostructure for the detection and inhibition of SARS-CoV-2 infections. Our experiments clearly demonstrate the advantage of using pattern-matching and multivalent interactions for increasing binding avidity between the DNA Net-aptamers and the SARS-CoV-2 virion. Characterizing the solution dynamics of these nets at single molecule using cryo-EM imaging, and MD simulations on the full DNA Net and whole SARS-CoV-2 virus are worth a future investigation, which can potentially provide advanced structural insights to the physical properties of the DNA Net in the solution and can help in better understanding the behavior of different Nets. Furthermore, following the SARS-CoV-2 detection strategy, our multi-layer DNA Net design can be adapted to diagnose and/or combat other viruses that possess envelope glycoproteins like the SARS-CoV-2 spike timers and pose a severe risk to the human health in foreseeable future.

## Supplementary Information

Materials, Methods, Figs. 1 to 17, Table 1, Videos 1 and 2, and References.

## Acknowledgements

This work was supported in part by grants from the NSF (CBET RAPID 20-27778) to B.T.C. and X.W., and NIH/NIAAA (AA029348) to X.W., B.T.C., and W.H. The authors thank Dr. Longping Victor Tse at the University of North Carolina and Drs. Jinghe Huang, Lu Lu and Fan Wu at Fudan University for helpful discussions about SARS-CoV-2 inhibition assays, Dr. Yongjun Guan at Antibody Biopharm Inc. for the anti-SARS-CoV-2 N protein monoclonal antibody, and Drs. Joel Baines and Claire Birkenheuer at Louisiana State University for VERO cell line.

## Author Contributions

N.C. performed all the sensing assays. S.R. crunched the DNA sequences. Y.X. and N.C. performed agarose gel electrophoresis (AGE) and AFM imaging to characterize DNA net structures. A.D. performed docking and MD simulation of aptamer-spike interaction. L.Z. performed coarse-grained simulations. X.L. and S.Y performed SPR assays. Y.X. performed confocal imaging assays. N.C. performed AGE to test the DNA net stability at different storage conditions and in saliva. N.M., T.Z. and W.H. performed the viral inhibition assays. B.T.C. provided advice on sensing instrument selection. X.W. conceived and supervised the study. N.C., YX, and X.W. led the preparation of the manuscript with inputs from other authors.

## Competing Financial Interests statement

A U.S. provisional patent has been filed in November 2020 based on part of the study reported in this manuscript.

